# Linear Embodied Saliency: a Model of Full-Body Kinematics-based Visual Attention

**DOI:** 10.1101/2020.02.08.938514

**Authors:** William W. Abbott, J. Alex Harston, A. Aldo Faisal

## Abstract

Gaze behaviour and motor actions are fundamentally interlinked in both a spatial and temporal manner. However, the vast majority of gaze behaviour research has focused to date on reductionist head-fixed screen viewing experiments and ignored the motor aspect of visuomotor behaviour, thereby neglecting a critical component of the perception-action loop. We address this with an experimental design to capture, rather than constrain, the full range of simultaneous gaze and motor behaviour in a range of natural daily life tasks. Through building autoregressive models and applying these to our novel datasets we find that beyond simple static regions of interest, we can predict visual attention shifts from freely-moving first person body kinematics, through explaining gaze dynamics in the context of body dynamics, on the timescale of freely moving interactive behaviour in individuals, expanding our understanding of natural visuomotor behaviour.

## Introduction

Eye movements are tightly linked to our behaviour and cognition through the allocation of overt and covert attention (Clark, 1998; M. F. Land & Furneaux, 1997), and also provide a direct proxy of attention, in that the fovea of the retina itself directly constrains high resolution information to a very small region of central vision (1-2°) (Hendrickson & Yuodelis, 1984). A key question is what guides our overt attention in space (i.e. fixation locations) in natural environments, and to understand how this is related to the task or action currently being performed. Central to this issue is what has become known as the saliency argument, one which debates how much the control of eye movements is ‘bottom-up’, based on visual conspicuity, or ‘top-down’, based on cognitive influences and task context (Schutz et al., 2011; Tatler et al., 2011). Visual conspicuity, or saliency, refers to locations in an image that “pop-out” from their background due to high contrast in image features such as colour, intensity or orientation. These features can be analysed at different spatial scales, and integrated to create a conspicuity map for each of these features, closely resembling the parallel processing carried out in the primary visual cortex (Carandini et al., 2005). These maps for each feature are normalised and combined into a salience map (Faisal et al., 1998; Itti et al., 1998; Koch & Ullman, 1985). Such models can then predict fixations at the most salient feature points of the image in a winner-takes-all fashion, with subsequent local inhibition of return applied to prevent immediate re-fixation. Bottom-up models have received huge attention both because they reflect our neurophysiological understanding of processing pathways and because they are computationally tractable, and such models can be applied and tested to novel images, with modern bottom-up saliency models correctly predicting the majority of ground-truth human fixations in head-fixed free-viewing of static images (between 0.53-0.88 ROC AUC (Area Under the Receiver-Operator Curve), chance level = 0.5 (Betz et al., 2010; Bylinskii et al., 2015)). The large majority of saliency research has to date focussed more on bottom-up image features and less so on task demands. Recent developments in the saliency field include significant improvements to methodology through the use of deep learning architectures and incorporation of top-down information (Borji et al., 10/2013, 2011; Kummerer et al., 2017).

Computational modelling of bottom-up visual saliency in this way has led to multiple competing computational saliency models (Bylinskii et al., 2015; Itti et al., 1998; Kümmerer et al., 2015), which provide direct predictions that can be applied and tested in static head-fixed 2D-screen-viewing experiments, with varying levels of performance. These models simply capture spatial allocation of attention integrated over time, rather than dynamic movements. Whilst these models work well for static scene-viewing experiments, natural gaze distributions differ largely in real world settings due to dynamic object movement in scenes, movement of individuals relative to their surroundings and the effect of task context.

The predictive power of these bottom-up saliency models is observed with static 2D stimuli and head-fixed eye-tracking, where subjects are not given any task, and freely view the image whilst their head is mounted rigidly with respect to a monitor. It has been shown however that any predictive power achieved by these bottom-up models becomes insignificant if the subject is given a specific visual task, such as search (Einhauser et al., 2008; Henderson et al., 2007). These bottom-up models have also been tested outside of the lab, in more ecologically valid experiments, involving free body and head movements during outdoor walking. Mounting evidence points to the significant impact of top-down influences in addition to bottom-up image features (Einhauser et al., 2008; Onat et al., 2014; Tatler et al., 2011).

Recent work has highlighted the importance of accounting for such top-down effects in saliency, but such experiments also comprise static viewing conditions (Henderson et al., 2018; Peacock et al., 2019) and suggest a combined role wherein object features guide fixations with low-level saliency guiding prioritisation of sequential fixations. However, simply introducing dynamic video stimuli to head-fixed viewers (macaques and humans) demonstrates that low-level saliency models cannot account for the variance of natural gaze behaviour (Shepherd et al., 2010), even before factoring in the considerations of free movements into natural visuomotor behaviour. Low-level visual saliency clearly holds predictive power, yet there are a number of arguments that further reduce the significance of this already modest predictivity. Firstly, simple known gaze behaviour biases, such as the tendency to fixate at the centre of the screen (central fixation bias) can achieve comparable prediction rates to many static saliency models (Kienzle et al., 2009; Tatler, 2007; Tatler & Vincent, 2009). Central fixation bias has also been observed in ecologically relevant experiments where central fixation bias endures in freely behaving subjects walking outside (Foulsham et al., 2011; Hart et al., 2009).

Secondly, much of the predictive power of these models can be removed by altering the task (Einhäuser et al., 2008; Foulsham and Underwood, 2008; Henderson et al., 2007). This is perhaps not surprising, as task is bound to affect where we look, allowing us to obtain the specific information we require. Whilst low-level feature maps clearly do influence attention (Treisman & Gelade, 1980) and are grounded in neurophysiology (being extracted in V1 and representing visually descriptive regions of an image (Carandini et al., 2005)), other forces act to guide attention as context and goals become more complex. Different control loops have been shown to contribute to the assignment of our overt attention alongside simple bottom-up saliency (Schutz et al., 2011), including motor plans, a value function tailored to the ongoing task, object recognition and their probable location (Peacock et al., 2019). Such integration of top-down and bottom-up signals has been proposed to occur at higher levels in the visual hierarchy in so-called ‘priority maps’ for which neurophysiological studies are beginning to provide evidence (Schutz et al., 2011). In addition, Follet et al. compared empirical saliency maps of focal and ambient fixations based on their automated classification and found that focal fixations are more associated with visual features than ambient ones, providing further evidence for a distinction between bottom-up and top-down signals (Follet et al., 2011).

Since the pioneering work of (M. F. Land & Lee, 1994), who took eye-tracking studies out of the lab and into the wild, it has been established that eye-movements are tightly linked, spatially and temporally, with motor actions in the wild. ((Epelboim et al., 1995; M. M. Hayhoe et al., 2003; M. F. Land & Furneaux, 1997; M. Land et al., 1999; Patla & Vickers, 1997; Pelz & Canosa, 2001; Schutz et al., 2011; Tatler et al., 2011). However, in current computational models, motor information is not included directly as an input. There is a growing body of literature that attempts to model the task control of eye-movements by breaking behaviour down into subtasks and their prescribed visuomotor behaviour (M. Hayhoe & Ballard, 2014; Henderson, 2017). Using concepts of reward, uncertainty and task structural constraints, the sub-task sequence is predicted and thus the associated visuomotor behaviour can be predicted. These approaches provide important theoretical frameworks, but in practice require significant abstraction, specification of tractable reward functions and painstaking hand annotation of sub-tasks. As such, they can only be applied in simple defined scenarios that are either virtual and thus pre-defined, or simple enough to enable straightforward state estimation. Thus these models, for now, have limited use and the level of abstraction means they cannot explicitly explain eye movement dynamics.

When performing motor actions, we fixate primarily on objects or regions that are directly relevant for task completion – even if this is a location on the featureless kitchen worktop that we plan to place a cup, or the blank wall we expect to observe a squash ball to bounce on (Shafti et al., 2019; Tatler et al., 2011). Indeed, the use of gaze information as a control signal can enable (Shafti et al., 2019) human-robot interactions or improve reinforcement learning agents (Makrigiorgos et al., 2019). In addition, the temporal arrangements of fixations are directly time locked to the actions they relate to, and thus information is acquired in a “just in time basis” (Ballard et al., 1995). This spatial and temporal relationship between overt attention and action has been shown to be highly consistent across subjects, suggesting a common mechanism (Land et al., 1999).

Until a few years ago, gaze behaviour research was restricted to laboratory settings due to the complexity and non-portability of eye tracking equipment. However, previous work was restricted to static scene viewing, which only represents a very small subset of gaze behaviour. Technological improvements (Abbott & Faisal, 2012; Schneider et al., 2009) have led to more affordable head-mounted eye trackers, facilitating real-world behavioural study.

This change in paradigm has brought to light many insights into gaze behaviour in more ecologically relevant contexts, such as when its allocation is based on the coordinated movement of the body and the eyes. Work in this area has further demonstrated that the control of where we look is based, overwhelmingly so, on the location of information required by ensuing action sub-goals (Epelboim et al., 1995; Hayhoe et al., 2003; Land and Furneaux, 1997; Land et al., 1999; Patla and Vickers, 1997; Pelz and Canosa, 2001, Schütz et al., 2011; Tatler et al., 2011).

If we are to understand from the mounting body of evidence described above that gaze behaviour exists as part of an embodied system (Sprague et al., 2007), then to truly understand natural gaze behaviour we must design experimental methods to capture and incorporate this embodiment, rather than remove it. One significant boundary for modelling the link between perception and action is the lack of high-resolution body movement information. Given our understanding that eye-movements are embedded in a rich visuomotor repertoire, models to predict eye-movements should be data-driven, not only from a visual saliency perspective but from an embodied perspective.

We use the term ‘embodied’ here to imply placing perception into the context and the actions of the body, i.e. movement behaviour. Towards this, we present an embodied methodology here to automatically capture, rather than constrain, sensory inputs and motor outputs in natural behaviour. Sensory inputs are recorded using a head mounted eye-tracker, scene camera and microphone, whilst simultaneously recording skeletal motor outputs through motion tracking 51 degrees-of-freedom (DOF) in the body and 40 DOF in the hands.

Considering we make up to 5 eye movements per second, attempting to interpret these dynamics with action sub-tasks defined on a timescale of about 3 seconds per action confounds timescales and overlooks information contained in the natural dynamics. Thus, objective and scalable (automated) methodologies to capture and quantify unconstrained behaviour are required to make such a problem tractable. All our tracking equipment was wearable and markerless, therefore it allowed unconstrained behavioural monitoring “in the wild” without concerns for occlusions or other technical issues. This allows us to test the simple hypothesis of embodied saliency: can eye movements be predicted directly from the dynamics of the body?

## Methods

### Experimental Setup

We used a portable eye-tracker (SMI Eye Tracking Glasses 30Hz, Sensomotoric Instruments, Teltow Germany) in combination with a portable full body motion capture suit, measuring 51 degrees of freedom (DOF) from the body using 17 inertial measurement units (IGS-180 Animazoo, Brighton UK), 22 DOF from the right hand (Cyberglove 1) and 18 from the left (Cyberglove 3, Cyberglove Systems, San Diego, California, USA). All tracking equipment is markerless and thus allows extensive behavioural monitoring “in-the-wild”. Experiments were filmed with a static video camera, as well as with the integrated egovideo from the eye tracker camera. Online annotation was additionally made by one of the experimenters to separate tasks from sub-tasks.

#### Body tracking

The movement from the entire body (excluding fingers) was recorded using an IGS-180 motion capture suit (Animazoo UK Ltd, Brighton, UK) with integrated sensor nodes positioned to track key degrees of freedom in the body. Each sensor is a 9-axis inertial measurement unit (IMU) composed of a 3 axis accelerometer, 3 axis gyroscope and 3 axis magnetometer. The body sensor network is composed of 17 sensor nodes placed on each body segment as illustrated in Figure 1A. Sensor fusion was applied on board, with the suit streaming 51 degrees of freedom at 60Hz. Calibration of the motion capture suit was performed by a simple routine provided by the manufacturer as part of the control software.

**Figure 1:**
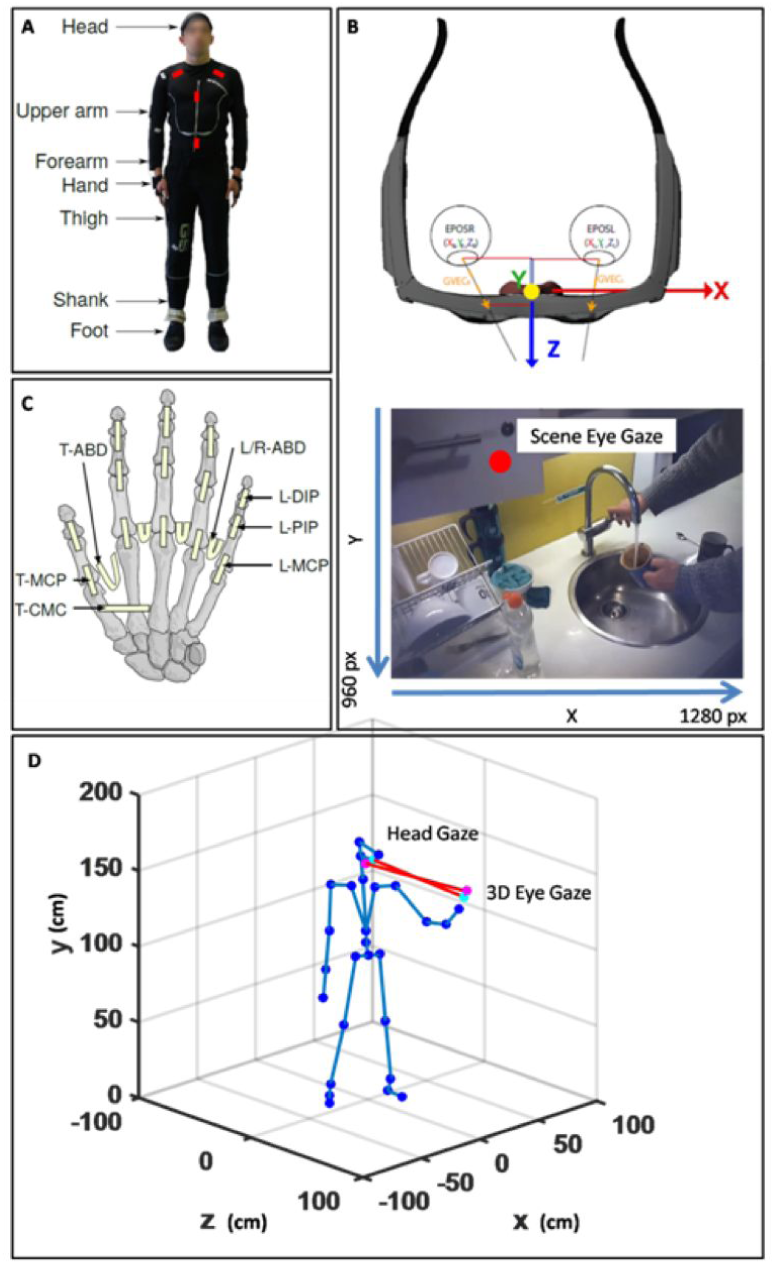
Eye movement data was collected using SMI Eye tracking glasses which record binocular eye-movements at 30Hz, synchronised and aligned to an egovideo feed (from the wearer’s point of view). Software was written in C++ using the SMI API to communicate with the eye-tracking server. Calibration checkpoints were made to check calibration periodically due to the length of the experiment, at 5 minute intervals.

#### Eye tracking

Eye movement data was collected using a pair of head-mounted SMI eye-tracking glasses (Fig 1B), allowing simultaneous recording of egovideo and pupils, allowing for full reconstruction. Custom software was written in C++ to interface the SMI API with the eye-tracking server. Calibration was performed according to standard SMI procedures at periodic intervals (every 5 minutes).

#### Hand Tracking

The data from the subjects’ left and right hand was recorded using a CyberGlove I and III respectively (CyberGlove LLC, San Diego, CA, USA). Note that the right hand glove also measures the movement of the distal interphalangeal (DIP) joints, while the left hand does not. The hand movement was recorded at 140Hz for the left and 90Hz for the right hand, and converted using an 8-bit analogue-to-digital converter (ADC). Calibration was performed as a two stage-process: an initial calibration using software provided by the manufacturer and subsequent refinement by manually adjusting sensor gains and offsets while comparing the subject’s hand with a visual rendering of the hand. The recorded data was streamed to a laptop and saved along with a time-stamp obtained from the system clock.

#### Scene Recording

The working area of the subject was monitored with a tripod mounted video recorder placed to capture the full scene. In addition the experiment was filmed from the subject’s perspective with the integrated scene camera on the head mounted eye-tracker.

### Ethomics Experiments

The experiment comprises 3 natural scenarios of daily life: breakfast time, evening chores and navigation. High level instructions are given such as “please lay the table for two people” or “please go ahead and have breakfast”. In the kitchen, subjects conducted high-level activities such as laying the table for breakfast, making tea, preparing and eating breakfast, and clearing up afterwards. The bedroom scenario involves evening chores and leisure activities including sweeping the floor, making the bed, packing their bags, playing games and socialising. One experimenter interacted with the subject and guided them through the tasks where absolutely necessary and interacted in tasks requiring a partner e.g. having breakfast together or playing games. The second experimenter observes and performs the online annotation.

### Data integration and curation

To collect such rich time-locked data, these collection modalities must run on distributed systems simultaneously with the ability to synchronise and integrate data post hoc. Time synchronisation was achieved with two approaches, primarily relying on network time protocol (NTP), which allows different systems to poll a universal clock to maintain a common time reference. A secondary backup system involved manual synchronisation via button press, injecting timestamp markers into the respective recording systems. To manage the data, a database structure was created to associate and link files to subjects, scenario and recording modality. This database structure was used to extract all data associated with a subject and scenario, to synchronise the modalities and finally to build a data structure amenable to the analysis with requisite pre-processing. Once synchronised, the suit and glove data were sampled at the eye-tracker’s ego video frame rate (30Hz) to maintain data integrity.

### Modelling dynamics

To test the idea of Embodied Saliency, which reflects that movement behaviour of the body predicts eye movements, we must take an approach of identifying mapping from the body joint angles (51 degrees of freedom (DOF)) to the eye gaze (2 DOF). Three regressive models are used: gaze autoregression, gaze autoregression with body joint angles as exogenous inputs, and pure linear regression from body joints to gaze. Figure 4 gives an overview of these models. The number of delays used in the regression was optimised by a parameter sweep. Regression was made to training data using the Moore-Penrose pseudo inverse (Moore, 1920). For the autoregressive models, training was performed in open loop (Figure 5) with the true current state of the eye position and the body joints being mapped to the next eye position. This was then tested both in open loop, again with known current eye position and in closed loop multistep ahead performance (Figure 5), where the estimated current state of eye-position was used rather than the actual current state.

**Figure 2:**
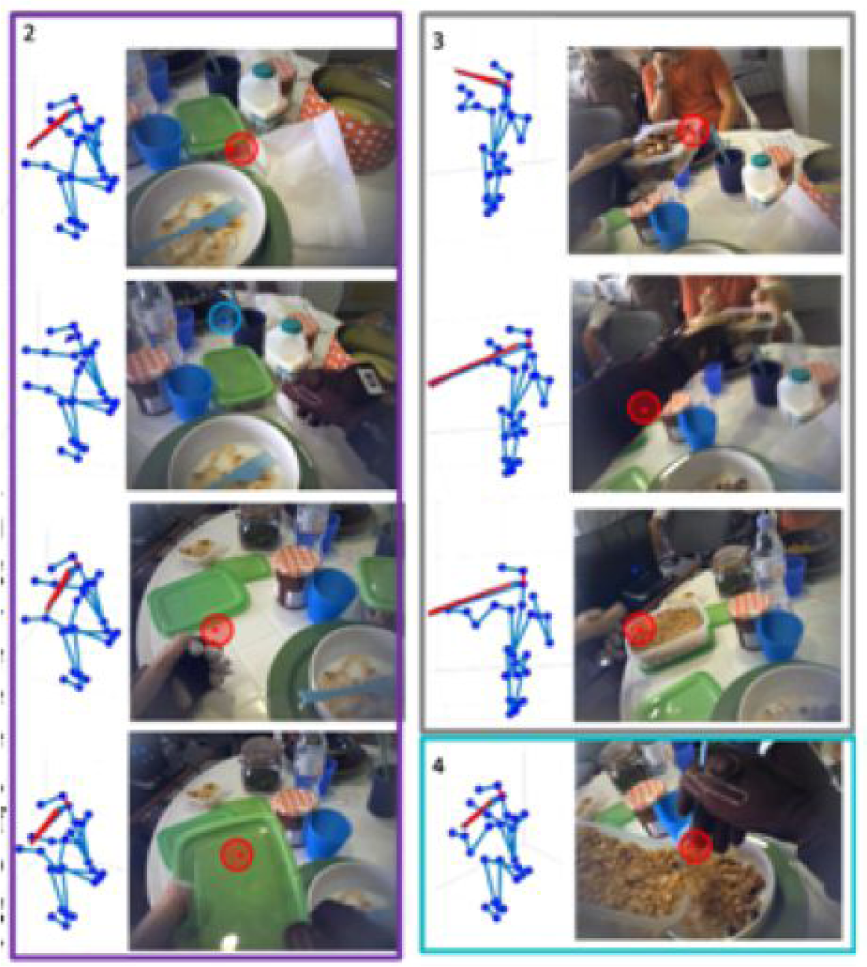
Example of full kinematics with 3D gaze endpoint reconstruction. The subject here is performing an activity of daily living - in this case, preparing and eating breakfast. Overview of the sensor suite and the corresponding coordinate systems. The suit and glove sensor placement is labelled in Figure 1A and 1C respectively. The gaze tracker coordinate system is shown in Figure 1B, which includes the 3D gaze vectors and the gaze with respect to the scene video. The integrated suit and gaze skeleton model was used with the joint angles from the suit to create a 3D model of the body first. Using the head position and absolute rotation the gaze vectors are rotated and translated to be relative to the head (Figure 1). 3D gaze was then estimated based on the GT3D system (Abbott & Faisal, 2012). In Figure 1 the blue line protruding from the head represents the head gaze orientation and the red lines represent the gaze vectors for each eye, which terminate at their closest point of intersection. The mid-point of these positions is the 3D gaze estimation.

**Figure 3:**
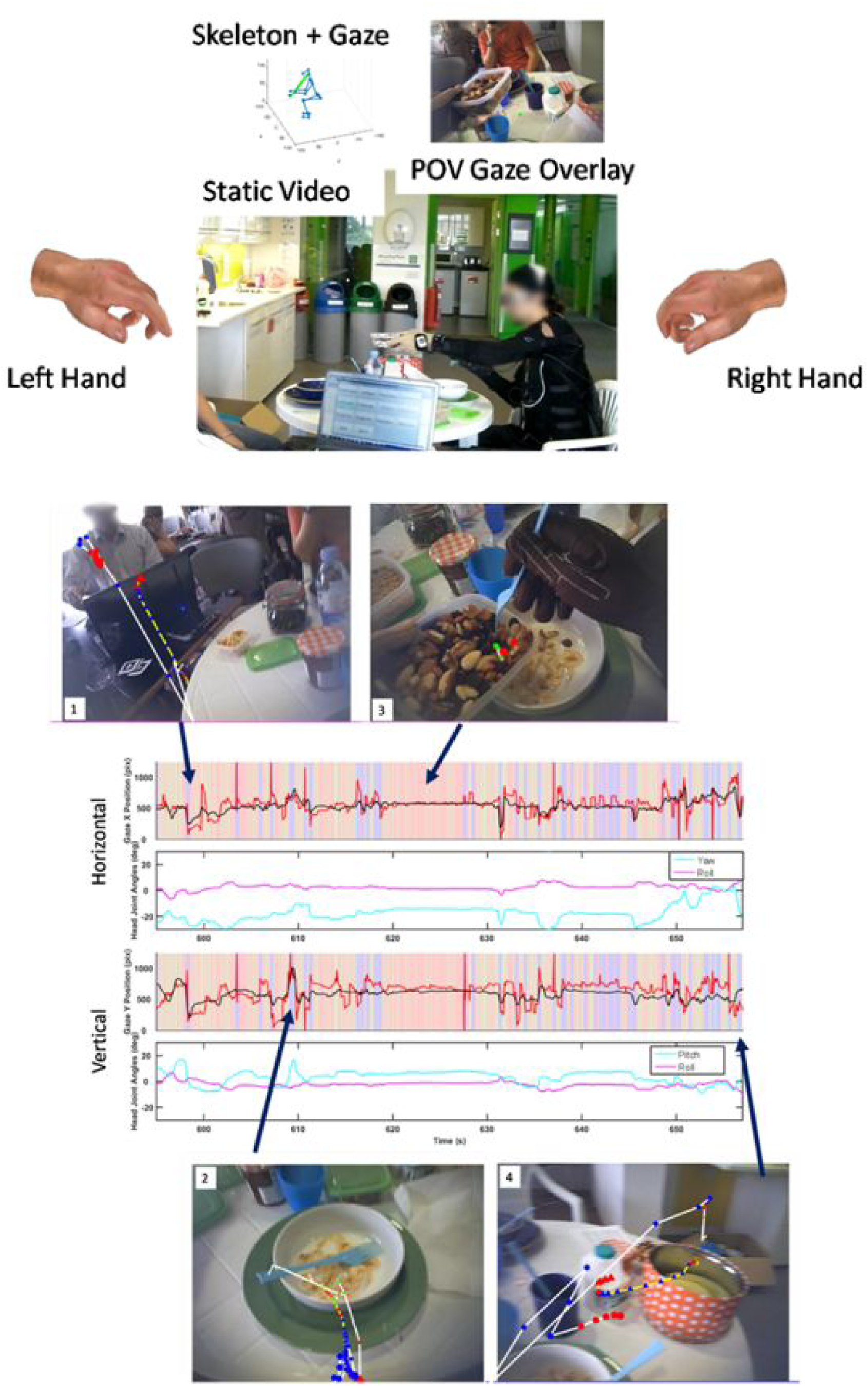
System overview of full scene reconstruction, complete with full hand reconstruction, egovideo, full skeleton kinematics and gaze tracking with external scene cameras.

**Figure 4:**
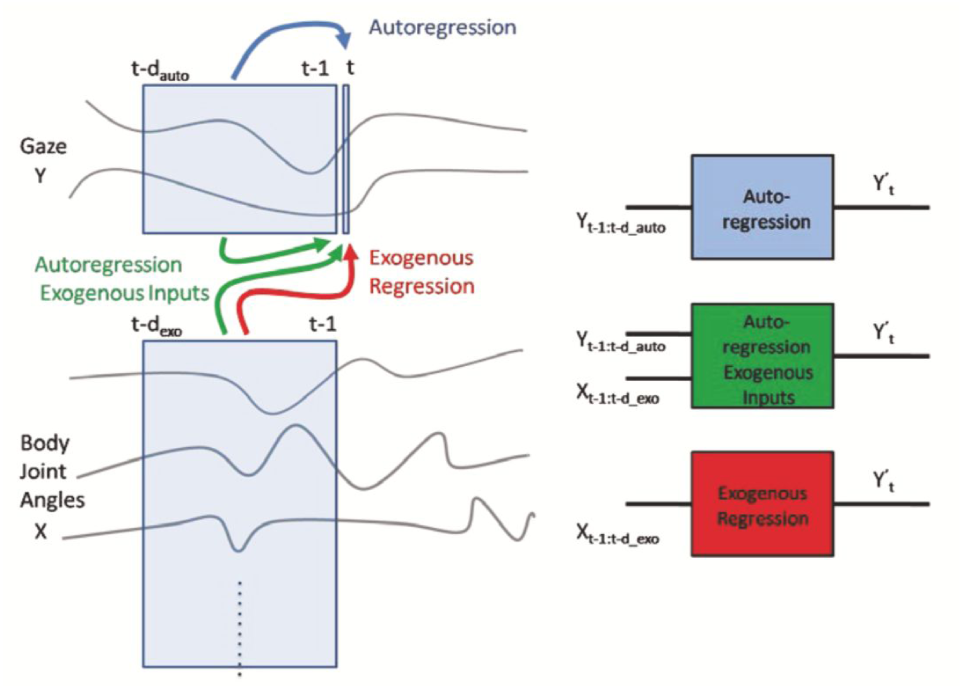
Linear regression models. Y denotes gaze position (pitch, yaw) and X denotes suit joint angles. Three linear models are used: Autoregression (blue), predicting future gaze based solely on past gaze. Autoregression with Exogenous Inputs (green), predicting future gaze based on past gaze and body joint angles. Exogenous regression (red), predicting future gaze based on past body joint angles. For each model the parameter that must be set is the number of delays (d_auto_ and d_exo_ depending on the model) used to predict gaze. Note: The “:” notation denotes all integers between, for example 1:5 means 1,2,3,4,5. Here t refers to frame number, so Y_t-1:t-d_ are the gaze positions for the previous d (delay) frames.

**Figure 5:**
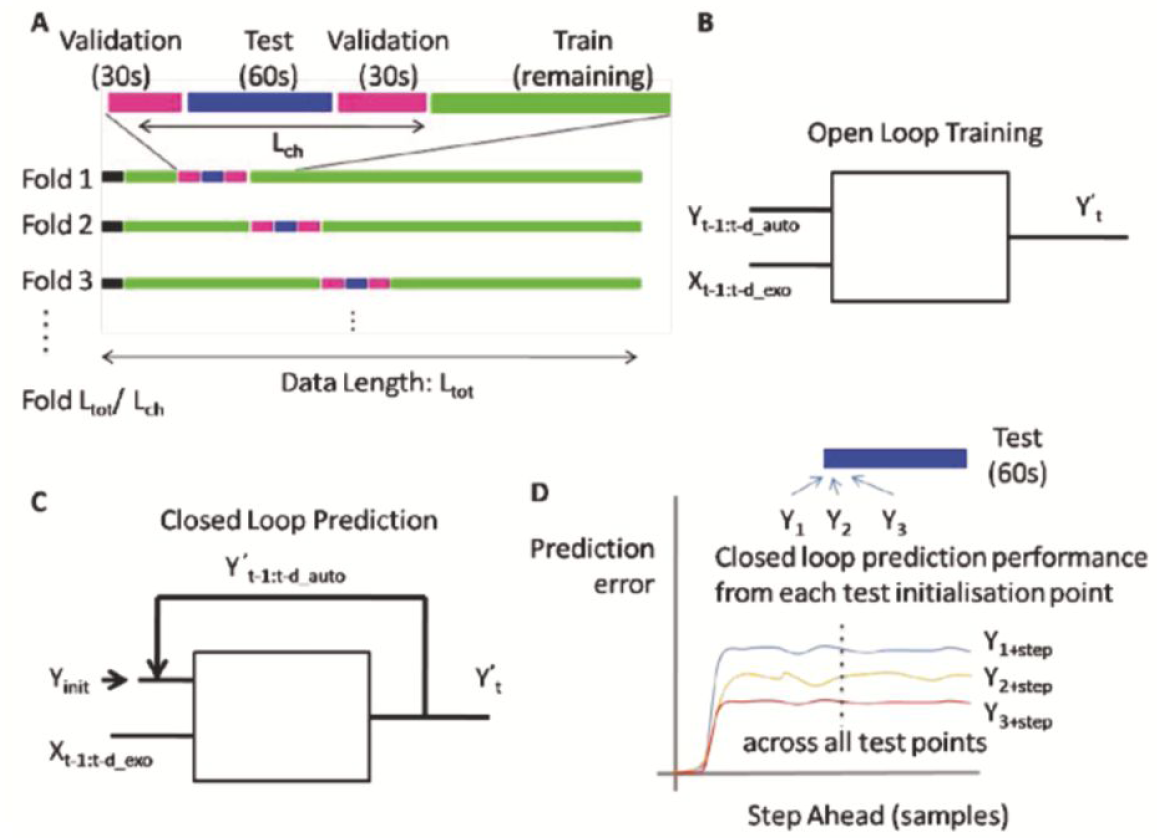
Training regime. A. Crossvalidation is performed by splitting data into training, validation and test datasets. Test data is flanked by validation data, providing separation between the test and training data. for each run on the same data set, the test chunk position is stepped through the data until all data has been used to train and test/validate. B. For autoregressive models, training is performed in open loop, meaning that the real past gaze data is available. Open loop performance comprises the prediction of the next state given the true current state. C. In closed loop mode the initial gaze position is known (Y_init_) for the first step prediction, after which the prediction is fed back to be used in subsequent step predictions. D. The prediction error is assessed by looking at the closed loop step ahead performance starting from each test data point location and predicting n steps ahead.

## Model Training & Validation

We take a batch based approach to training and testing. We split the data streams into training and test sections. Here it is important to consider the autocorrelation function to ensure the training and test data are sufficiently temporally separated. We assess the span of the temporal autocorrelation by taking the full width at half maximum (FWHM) of the autocorrelation function. The eye movement data has a vertical FWHM of 780ms and horizontal FWHM of 600ms. This suggests that the data is significantly correlated over intervals of around 1 second. After 5 seconds the correlation function becomes close to zero and thus it is safe to use blocks of training and test data that are at least 5 seconds apart. To be even more conservative, training and test data are separated by 30 seconds, to ensure no direct temporal correlations between training and test data. The data during blinks (detected by our eyetracker’s SMI BeGaze software) was removed from the training data. Cross-validation was performed by selecting a period of test data flanked by a minimum of 30s gap, with the remaining data used to train the model. For each round of cross-validation the training block (including flanking gaps) was moved to the next contiguous position at the end of the current block (Figure 5A). This regime allows the performance to be assessed at different positions of the time course showing where generalisation of the model is successful but also provides a measure of “similarity” or “predictability” of the data chunk.

The autoregressive models were run both in open loop (true previous gaze position known) and closed loop (predicted gazed feedback). Linear regression by comparison only has a straight input-output relationship (cf Figure 4). The open loop performance was calculated for each test data point, whilst for the closed loop performance the multi-step ahead performance was calculated, taking each test data point, providing the current real gaze position (as well as required lag values before) and then predicting each subsequent gaze position. Thus for subsequent steps the predicted gaze feedback (rather than true gaze) is used to predict the next steps. This multi-step ahead performance is then averaged across each starting position. The model performance is assessed on two metrics, the average (mean) absolute error (AAE) and the squared Pearson correlation coefficient (R^2^). The former assessing the actual error in the prediction and the latter capturing the linear correlation between the prediction and the actual gaze data, thus preventing non-stationary effects of drift in the sensors from shrouding performance. To provide a baseline comparison or “chance level” two methods are used: 1) Random Sampling: the gaze target estimate is made by sampling from a uniform distribution across the possible gaze range. 2) Central Fixation Bias: a normal distribution is fitted to the training data gaze distribution and then sampled from to predict the gaze location.

Assuming linear model dynamics with Gaussian noise ε we obtain the following Vector AutoRegressive model with eXogenous input (VARX) for predicting gazepoint *y* at time *t*, as defined in (Lütkepohl, 2005):

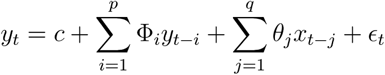

## Results

We set out to predict the direction of visual attention in head-centered coordinates with a set of time series models from body pose data alone. We used simple gaze autoregression, gaze autoregression with exogenous body input, and linear regression from body movements to gaze, as defined in the Methods section (Models: see Figure 4; Training: Figure 5). One critical measure is how good our models are at using data up to time t=0 to predict eye movements occurring at a future time point t=T. If our models are correct we should be able to predict the dynamics of future eye movements by rolling out (see Methods) next time step predictions of eye movements, feeding them back into the system up to our desired time point T. We executed this rollahead for a prediction horizon ranging from T=0 to T=10s. We present the results in terms of the average absolute error (AAE), which should increase with T (up to steady state error where the limits are set by boundary effects of eye movement range and central fixation bias) and R^2^, which with increasing T should decrease as unforeseeable future events will make eye movements unpredictable. We are focussing therein on the intermediate range of prediction horizons after the very immediate future (0-10ms), on the timescale before predictability of the world and task are limited by their own dynamics (approx 2s).

We focus here on the average rollahead prediction AAE and R^2^ (averaged across all subjects, across scenarios and across the length of each recording session for the three tested models). Simple linear regression from the dynamics of the body to predict gaze (“Exo”, red line, Figure 6) is relatively constant across prediction horizons (between 4-4.5 deg AAE and in terms of R^2^ 0.35-0.425). In contrast, autoregression (predicting gaze from gaze; see Figure 6 blue line) and our autoregressive with exogenous input model (see Figure 6, green) perform significantly better on very short time scales (<200ms) and in line with visuomotor reaction time (approx 150-200ms, (Hülsdünker et al., 2017). After 200ms the autoregressive prediction with exogenous inputs outperforms all other prediction models up to durations of a few seconds in terms of AAE and R^2^. Crucially all 3 model frameworks perform significantly better than the central fixation bias which is 200-300% higher in terms of AAE error. Moreover, central fixation bias predicts only state visual attention probability distributions, and no dynamic component of eye movements, unlike our models.

**Figure 6:**
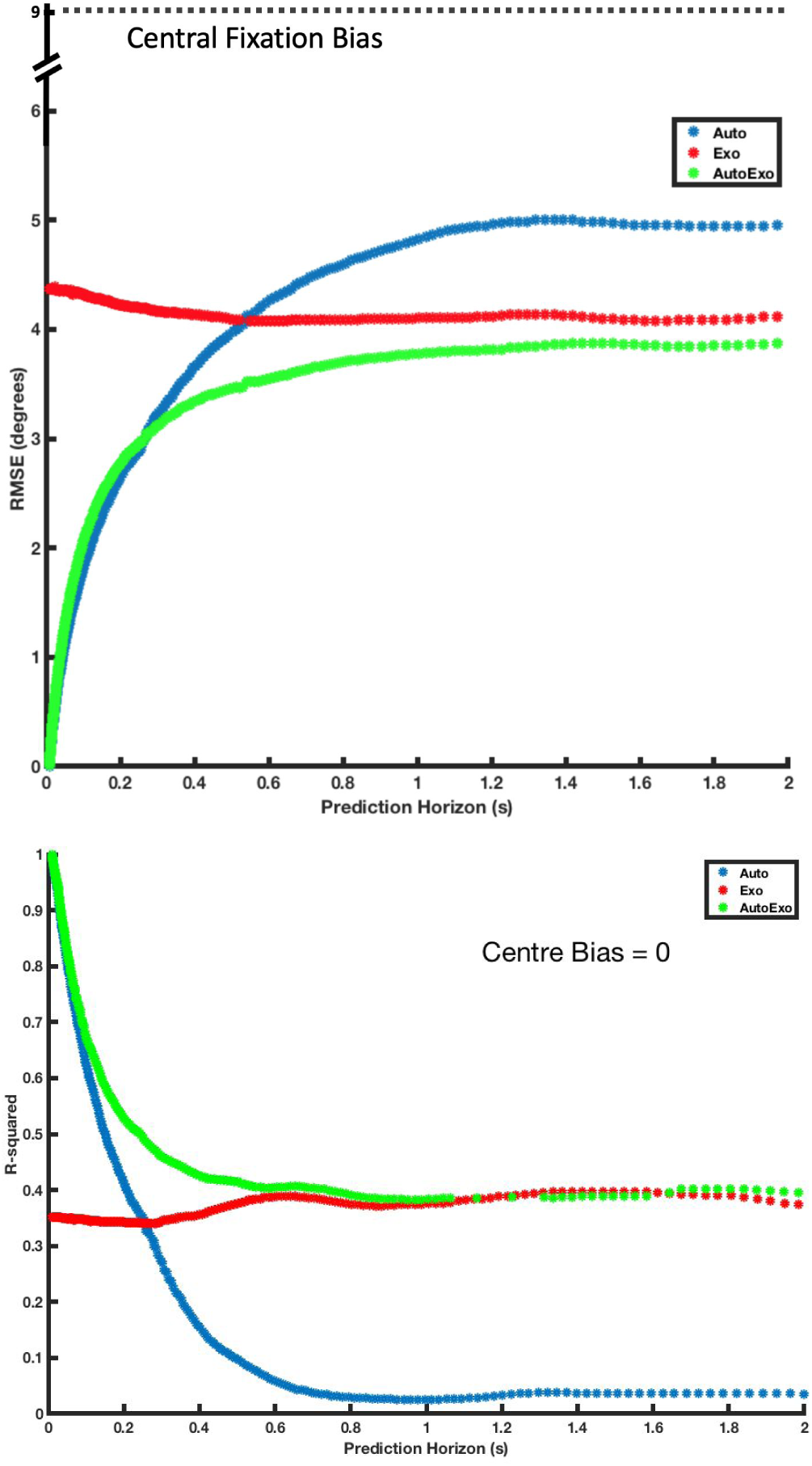
Averaged closed loop rollahead prediction for simple gaze autoregression (‘Auto’, blue), gaze autoregression with exogenous body input (‘AutoExo’, green) and linear regression from body to eye (‘Exo’, red).

We find that across different real-world scenarios, gaze autoregression with exogenous body input consistently outperforms central bias and random uniform sampling in terms of both AAE and R^2^ metrics (Figure 7). This is likely due to the combination of short-term dynamics with respect to the eye movements. There is also a nonsignificant difference between autoregression with exogenous body input and exogenous regression from body to eye across both bedroom and kitchen scenarios (Figure 7A, 7B) Different limb sets were also tested to investigate the different impacts of particular joint sets on gaze predictivity - we tested multiple possible combinations of limb joints to understand the relative impacts of different joints on prediction performance. All joint sets tested outperform centre bias and chance level with respect to both mean error and R-squared metrics. Joint sets that include head dynamics (‘Head, Spine and Hips’; ‘Just Head’; ‘Head and Arms’) outperform the joint sets that do not, indicating the impact of head dynamics on gaze in real-world behaviour.

**Figure 7:**
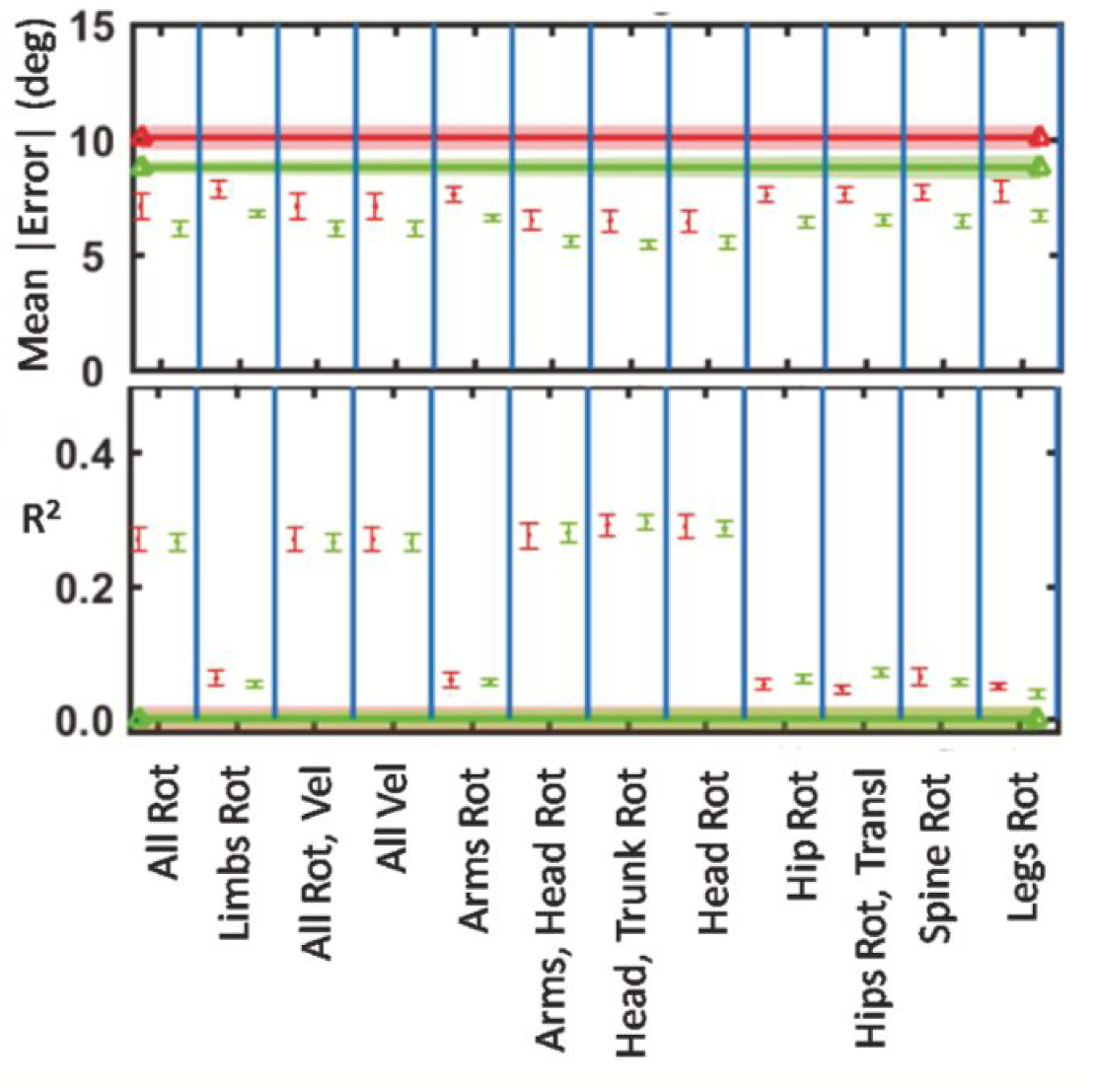
Performance overview. A. Comparison of mean absolute error for each regression method to predict eye gaze (cross is autoregression with exogenous suit inputs, dot is linear regression from suit inputs) in kitchen (red) and bedroom (green). All joint angles used, plotting mean and standard error cross validated and across all subjects. B. As for A, but comparing R^2^ (Pearson’s squared correlation coefficient). Here the linear regression from suit to eye gaze, pairwise, has a significantly better rollahead performance than the autoregression with exogenous suit inputs, (two tail paired t-test, kitchen p=0.03, bedroom p=0.02). The difference between settings is not significant. C. Comparison between linear regression (dot) from all suit sensors to gaze and two “chance” metrics. The first metric (CB) is the common saliency benchmark based on the central fixation bias (see methods). The second metric (rand) is based on the performance with gaze estimates sampled from a random uniform distribution. E and F compares the mean absolute error and R^2^ for linear regression with different combinations of joint inputs. ‘Rot’ here denotes joint rotation angle and ‘Vel’ denotes the joint’s angular velocity. For example All Rot, Vel uses the rotation angle for all joints concatenated with the rotation velocity for all joints. The straight lines plotted across the figure represent the central fixation chance level for each setting.

Different lag parameters were tested to optimise the models. (Figure 8). Multi-step ahead performance for the autoregression with exogenous input model plateaus at around 0.35 R^2^ (Fig 8A) for the horizontal component and 0.22 R^2^ for the vertical component. The RMSE stabilises at 170 pixels (8 degrees) in the horizontal and 210 pixels (10.1 degrees) in the vertical. This optimal linear prediction is achieved with 2 input delays in both autoregressive feedback and exogenous input components. The x-axis labels in Fig. 8B indicate the time lags from 1 single frame, up to 15 lags, as well as logarithmically spaced lags. Each frame is 33 ms apart from the next, so a 10 lag frame is approximately. 1/3 of a second prior.

**Figure 8:**
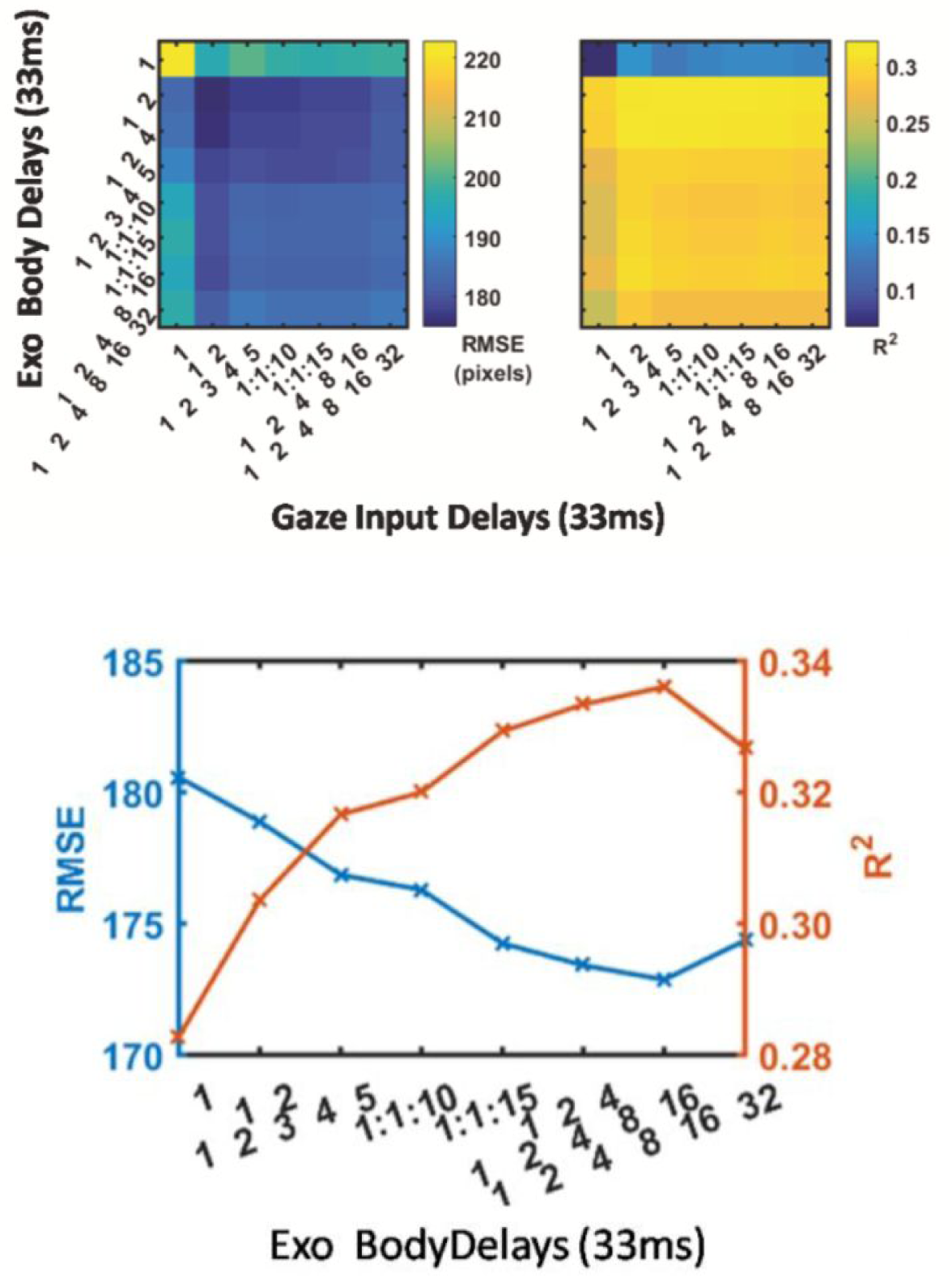
A. Autoregression with Exogenous Input model hyperparameter selection. Multi-step ahead performance plateaus at around 0.35 R^2^ for the horizontal component and 0.22 for the vertical component. The RMSE stabilises at 170 pixels (8 degrees) in the horizontal and 210 pixels (10.1 degrees) in the vertical. This optimal linear prediction is achieved with 2 input delays in both autoregressive feedback and exogenous input components. B. Autoregression Model Hyperparameter Selection C. Delay parameter optimisation in open loop for simple autoregression, comparing RMSE and R^2^ metrics for different input delays. The x-axis labels indicate the time lags from 1 single frame, up to 15 lags, as well as logarithmically spaced lags. Each frame is 33 ms apart from the next, so a 10 lag frame is approximately. 1/3 of a second prior.

## Discussion

Classically, allocation of gaze has been computationally modelled as bottom-up scene salience and qualitatively argued to depend on top-down ongoing task demands. We demonstrate here that rigorous computational modelling allows us to analyse and predict eye-movement dynamics - both the location and time course of attention - by relating them directly to body kinematics when acting in the wild. Thereby, the interpretation of gaze and attention is integrated, rather than disembodied, from the kinematics of motor behaviour. Previous data-driven methodologies have addressed eye-movement behaviour in the wild, but have focused on natural visual stimuli (Foulsham et al., 2011; Hart et al., 2009). Efforts have also been made to incorporate head movements (Einhäuser et al., 2007), however this is the only known study incorporating such comprehensive real-time body movement recordings. The embodied eye-movement methodology described herein has lead to the main concept presented here: the interaction of body posture and spatial allocation of overt attention, which we term ‘Embodied Saliency’.

We find that embodied saliency, as captured by our AutoExo model, best explains human eye movements at intermediate time scales (200ms-2000ms) while the autoregressive model operates better at very short delays (<200ms) where visual information intake is more predictable (e.g. it is faster than reaction and motor planning timescales).

Our approach of measuring embodied saliency attempts to capture the concept of interpreting the spatial allocation of gaze behaviour in terms of the pose of the body. Body pose dynamics carry both information about sensorimotor control requirements as well as information about the task and ultimately affordances that drive a subject’s behaviour and thus their visual attention.

We believe the embodied saliency concept is important when interpreting gaze behaviour in the wild, where gaze can no longer just be thought of as eye-movements in response to visual stimuli. Gaze allocation is thought of instead as a coordination of body, head and eye-movements. In addition, the actions of the body, which are recorded right down to every joint in the finger, must affect how this attention is allocated. This approach has previously been taken for recording head-movements and eye-movements simultaneously in the wild (Einhäuser et al., 2007).

We explore the idea that eye-movements can be predicted by the conformation and dynamics of the body. The body stance we take, and the way we move our head and limbs offers clues as to where we are likely to be looking. There is a large amount of previously-discussed literature that suggests this to be the case, linking eye-movement to action, but as discussed, this tends to be qualitative outside of reductionist lab settings, and whilst lab studies provide exciting justification, the findings need to be demonstrated in a similarly quantitative manner whilst maintaining the complexities of behaviour in the wild. By mapping here directly from full body kinematics to eye-movements, we show significant predictive power within body dynamics themselves.

We show here that body movements can be used to explain up to 40% of eye-movement variance with a simple linear model several seconds ahead in time, and thus it is important to investigate how the remaining variance can be attributed to other embodied interactions. The current linear model used is dominated by the synergistic effects of head and trunk movements in predicting eye-movements. While subtle effects are seen in other regions of the joint space such as the shoulders and hip position, their contribution do not significantly improve performance across the whole recording. This does not refute the claims of embodied saliency, but demonstrates, perhaps expectedly, that a linear model alone cannot fully capture the complex and multifaceted nature of these interactions. The model can now be extended and iteratively improved in the ways described below to explain more of the variance of eye-movement behaviour.

### Multiple Time Scales

Eye-movements occur on very different time scales, with rapid saccadic eye-movements and smooth slow eye-movements to completely static fixations. The same is true for different body parts which tend to act over considerably longer time-scales than the eye. For example, for shoulder movements the FWHM for shoulder elevation velocity is 600-800ms, compared to ∼100ms for the eyes. Thus when an object manipulation such as a reach is initiated, the gaze movement associated with it is long over by the time the end of the trajectory is reached. Therefore, using a constant linear filter to map from arm to eye movement is perhaps overly simplistic, particularly given that by the time the arm reaches the position that was fixated, the gaze is likely to have already moved on.

### Multiple Latencies

Another limitation of the current linear model is that the number of lags is a single optimisation parameter across all dimensions, rather than allowing different lags for different joints. Thus the optimal latency used to map from body joints to eye-movements is likely to be suited to leverage the dominant synergistic head movements but not be optimal for other joints. Further to this, there is likely useful information in proactive inputs, for example the hand position in 1 second may contain information about the current gaze position. It will be these components of the model that have important implications for prosthetic control, as the model can be inverted to predict future arm movements from the current gaze behaviour. These proactive planning modes must be separated however from the reactive proprioceptive modes, where the gaze is drawn to the current position of the hand, which introduces the next component.

### Multiple Modes

As has been discussed with regard to the head and trunk, the contribution to eye-movements can be both compensatory and synergistic. In the same vein eye movements associated with the hand can be both guiding and following - i.e the eye can lead the hand guiding the action, where in other modes the eye can be guided by arm position and proprioceptive inputs. Thus, future models must enable these different mechanisms to be captured and switch between them appropriately.

The fact that this central fixation bias appears to endure in natural behaviour counters arguments that suggest centre bias in static natural image viewing is an effect of photographer bias (Tseng et al., 2009). The explanation for central fixation bias enduring in natural behaviour could instead come from the fact that the head and indeed whole body movements are used in combination with the eyes to change gaze heading. While the eye gaze may arrive at the target first, as the head catches up the eyes return to central fixation and thus the amount of time eye-gaze needs to be held at extreme eccentricities is minimised. This would be beneficial for both energy constraints and minimising motor noise. This eccentricity minimisation bias, similar to the concept of orbital reserve, may then endure - even in head restrained screen viewing. Testing the hypothesis of embodied saliency, we show here that the head is a strong predictor of eye-movements, beyond central fixation bias itself.

It is often the experimental context and capabilities that shape emerging theories. As eye-movement studies have moved out of the lab and into the real world, the role of physical tasks on gaze patterns has become increasingly apparent. However, computational models are still driven predominantly by visual stimuli. Now, with the ability to record natural gaze and motor behaviour at an unprecedented resolution, all-encompassing data driven models in this manner can emerge, that work towards capturing the full complexity of behaviour through the integration of these multifaceted and simultaneous drivers of eye-movements, and thus attention.

## References

Abbott, W. W., & Faisal, A. a. (2012). Ultra-low-cost 3D gaze estimation: an intuitive high information throughput compliment to direct brain–machine interfaces. Journal of Neural Engineering, 9 (4), 046016.

Ballard, D. H., Hayhoe, M. M., & Pelz, J. B. (1995). Memory Representations in Natural Tasks. Journal of Cognitive Neuroscience, 7 (1), 66–80.

Betz, T., Kietzmann, T., Wilming, N., & Konig, P. (2010). Investigating task-dependent top-down effects on overt visual attention. Journal of Vision, 10 (3), 1–14.

Borji, A., Sihite, D. N., & Itti, L. (10/2013). What stands out in a scene? A study of human explicit saliency judgment. Vision Research, 91, 62–77.

Borji, A., Sihite, D. N., & Itti, L. (2011). Quantifying the relative influence of photographer bias and viewing strategy on scene viewing. Journal of Vision, 11 (11), 166–166.

Bylinskii, Z., Judd, T., Borji, A., Itti, L., Durand, F., Oliva, A., & Torralba, A. (2015). Mit saliency benchmark.

Carandini, M., Demb, J. B., Mante, V., Tolhurst, D. J., Dan, Y., Olshausen, B. A., Gallant, J. L., & Rust, N. C. (2005). Do we know what the early visual system does? The Journal of Neuroscience 25 (46), 10577–10597.

Clark, A. (1998). Being There: Putting Brain, Body, and World Together Again. MIT Press.

Einhäuser, W., Schumann, F., Bardins, S., Bartl, K., Böning, G., Schneider, E., & König, P. (2007). Human eye-head co-ordination in natural exploration. Network: Computation in Neural Systems, 18, 267–297.

Einhauser, W., Spain, M., & Perona, P. (2008). Objects predict fixations better than early saliency. Journal of Vision, 8 (14), 18–18.

Epelboim, J., Steinman, R. M., Kowler, E., Edwards, M., Pizlo, Z., Erkelens, C. J., & Collewijn, H. (1995). The function of visual search and memory in sequential looking tasks. Vision Research, 35 (23-24), 3401–3422.

Faisal, A. A., Fislage, M., Pomplun, M., Rae, R., & Ritter, H. (1998). Observation of human eye movements to simulate visual exploration of complex scenes. SFB Rep, 360, 1–34.

Follet, B., Le Meur, O., & Baccino, T. (2011). New insights into ambient and focal visual fixations using an automatic classification algorithm. I-Perception, 2 (6), 592–610.

Hayhoe, M., & Ballard, D. (2014). Modeling task control of eye movements. Current Biology: CB, 24 (13), R622–R628.

Hayhoe, M. M., Shrivastava, A., Mruczek, R., & Pelz, J. B. (2003). Visual memory and motor planning in a natural task. Journal of Vision, 3 (1), 6.

Henderson, J. M. (2017). Gaze Control as Prediction. Trends in Cognitive Sciences, 21 (1), 15–23.

Henderson, J. M., Brockmole, J. R., Castelhano, M. S., & Mack, M. (2007). Chapter 25 - Visual saliency does not account for eye movements during visual search in real-world scenes. Eye Movements (pp. 537 – III). Elsevier.

Henderson, J. M., Hayes, T. R., Rehrig, G., & Ferreira, F. (2018). Meaning Guides Attention during Real-World Scene Description. Scientific Reports, 8 (1), 13504.

Hendrickson, A. E., & Yuodelis, C. (1984). The morphological development of the human fovea. Ophthalmology, 91 (6), 603–612.

Hülsdünker, T., Strüder, H. K., & Mierau, A. (2017). Visual Motion Processing Subserves Faster Visuomotor Reaction in Badminton Players. Medicine and Science in Sports and Exercise, 49 (6), 1097–1110.

Itti, L., Koch, C., & Niebur, E. (1998). A model of saliency-based visual attention for rapid scene analysis. IEEE Transactions on Pattern Analysis and Machine Intelligence, 20 (11), 1254–1259.

Kienzle, W., Franz, M. O., Schölkopf, B., & Wichmann, F. A. (2009). Center-surround patterns emerge as optimal predictors for human saccade targets. Journal of Vision, 9 (5), 7.1–15.

Koch, C., & Ullman, S. (1985). Shifts in selective visual attention: towards the underlying neural circuitry. Human Neurobiology, 4 (4), 219–227.

Kümmerer, M., Wallis, T. S. A., & Bethge, M. (2015). Information-theoretic model comparison unifies saliency metrics. Proceedings of the National Academy of Sciences, 112 (52), 16054–16059.

Kummerer, M., Wallis, T. S. A., Gatys, L. A., & Bethge, M. (2017). Understanding Low- and High-Level Contributions to Fixation Prediction. Proceedings / IEEE International Conference on Computer Vision. IEEE International Conference on Computer Vision, 4799–4808.

Land, M. F., & Furneaux, S. (1997). The knowledge base of the oculomotor system. Philosophical Transactions of the Royal Society of London. Series B, Biological Sciences, 352 (1358), 1231–1239.

Land, M. F., & Lee, D. N. (1994). Where we look when we steer. Nature, 369 (6483), 742–744.

Land, M., Mennie, N., & Rusted, J. (1999). The roles of vision and eye movements in the control of activities of daily living. Perception, 28 (11), 1311–1328.

Lütkepohl, H. (2005). New introduction to multiple time series analysis. New York : Springer.

Makrigiorgos, A., Shafti, A., Harston, A., Gerard, J., & Aldo Faisal, A. (2019). Human Visual Attention Prediction Boosts Learning & Performance of Autonomous Driving Agents. In arXiv [cs.CV]. arXiv. http://arxiv.org/abs/1909.05003

Onat, S., Açık, A., Schumann, F., & König, P. (2014). The contributions of image content and behavioral relevancy to overt attention. PloS One, 9 (4), e93254.

Patla, A. E., & Vickers, J. N. (1997). Where and when do we look as we approach and step over an obstacle in the travel path? Neuroreport, 8 (17), 3661–3665.

Peacock, C. E., Hayes, T. R., & Henderson, J. M. (2019). Meaning guides attention during scene viewing, even when it is irrelevant. Attention, Perception & Psychophysics, 81 (1), 20–34.

Pelz, J. B., & Canosa, R. (2001). Oculomotor behavior and perceptual strategies in complex tasks. Vision Research, 41 (25-26), 3587–3596.

Schneider, E., Villgrattner, T., Vockeroth, J., Bartl, K., Kohlbecher, S., Bardins, S., Ulbrich, H., & Brandt, T. (2009). EyeSeeCam: an eye movement-driven head camera for the examination of natural visual exploration. Annals of the New York Academy of Sciences, 1164, 461–467.

Schutz, A. C., Braun, D. I., & Gegenfurtner, K. R. (2011). Eye movements and perception: A selective review. Journal of Vision, 11 (5), 9–9.

Shafti, A., Orlov, P., & Aldo Faisal, A. (2019). Gaze-based, Context-aware Robotic System for Assisted Reaching and Grasping. 2019 International Conference on Robotics and Automation (ICRA), 863–869.

Shepherd, S. V., Steckenfinger, S. A., Hasson, U., & Ghazanfar, A. A. (2010). Human-monkey gaze correlations reveal convergent and divergent patterns of movie viewing. Current Biology: CB, 20 (7), 649–656.

Sprague, N., Ballard, D., & Robinson, A. (2007). Modeling embodied visual behaviors. ACM Transactions on Applied Perception, 4 (2), 11 – es.

Tatler, B. W. (2007). The central fixation bias in scene viewing: Selecting an optimal viewing position independently of motor biases and image feature distributions. Journal of Vision, 7 (14), 4.

Tatler, B. W., Hayhoe, M. M., Land, M. F., & Ballard, D. H. (2011). Eye guidance in natural vision: reinterpreting salience. Journal of Vision, 11 (5), 5.

Tatler, B. W., & Vincent, B. T. (2009). The prominence of behavioural biases in eye guidance. Visual Cognition, 17 (6-7), 1029–1054.

Treisman, A. M., & Gelade, G. (1980). A feature-integration theory of attention. Cognitive Psychology, 12 (1), 97–136.

